# A Dual Controllability Analysis of Influenza Virus-Host Protein-Protein Interaction Networks for Antiviral Drug Target Discovery

**DOI:** 10.1101/429712

**Authors:** Emily E. Ackerman, John F. Alcorn, Takeshi Hase, Jason E. Shoemaker

## Abstract

Host factors of influenza virus replication are often found in key topological positions within protein-protein interaction networks. This work explores how protein states can be manipulated through controllability analysis: the determination of the minimum manipulation needed to drive the cell system to any desired state. Here, we complete a two-part controllability analysis of two protein networks: a host network representing the healthy cell state and an influenza A virus-host network representing the infected cell state. This knowledge can be utilized to understand disease dynamics and isolate proteins for study as drug target candidates. Both topological and controllability analyses provide evidence of wide-reaching network effects stemming from the addition of viral-host protein interactions. Virus interacting and driver host proteins are significant both topologically and in controllability, therefore playing important roles in cell behavior during infection. 24 proteins are identified as holding regulatory roles specific to the infected cell by measures of topology, controllability, and functional role. These proteins are recommended for further study as potential antiviral drug targets.

**Importance:** Seasonal outbreaks of influenza A virus are a major cause of illness and death around the world each year, with a constant threat of pandemic infection. Even so, the FDA has only approved four treatments, two of which are unsuited for at risk groups such as children and those with breathing complications. This research aims to increase the efficiency of antiviral drug target discovery using existing protein-protein interaction data and network analysis methods. Controllability analyses identify key regulating host factors of the infected cell’s progression, findings which are supported by biological context. These results are beneficial to future studies of influenza virus, both experimental and computational.

## Introduction

The development of computational methods to identify key host factors that allow viruses to interrupt and control healthy cell functions will greatly aid in the prediction of novel anti-viral drug targets^1^. Traditional systems biology approaches to understanding cell dynamics during infection include the creation of detailed kinetic models for intercellular signaling pathways. While these models are advantageous in understanding the disease state in a quantitative way, they require experimentally-derived or estimated parameters and training data^2–4^, without which complications can arise and an accurate model can quickly become unattainable. Further, modeling studies are often limited to specific pathways which fails to consider the total cellular environment as an interdependent system.

Alternatively, network analysis methods applied to protein-protein interaction (PPI) data have been used to model cell-wide systemic changes associated with disease, changes in cell function, or cell fate^5^. This strategy provides a holistic understanding of system behavior by viewing proteins as interdependent states, regardless of specific interaction mechanisms, and allows for the exploration of cell level relationships. The field of network theory is well established; basic network metrics like degree and betweenness^6^ have repeatedly revealed the importance of specific proteins within biological processes that cannot be found from traditional modeling approaches^7–11^. Disease networks have identified genes involved with cancer^12–15^, demonstrated that the genes responsible for similar diseases are likely to interact with each other^16,17^, and predicted novel drug targets^18,19^.

There is precedent for network studies of many common viruses including hepatitis C^20,21^, SARS^16,22^, HIV^22–26^, and influenza virus^16,27–30^. Past work studying the effects of influenza virus in PPI networks has focused on identifying host factors involved in virus replication and improving the prediction of drug targets, but ends with an analysis of basic topological measurements. While this provides a general overview of the state of the network, it is a static snapshot of the cell and, therefore, fails to capture the dynamic nature of the cell. Therefore, the next logical step in analyzing biological networks lies in understanding how these dynamic systems can be manipulated and exploited to manage biological properties.

In classic control theory, controllability is the idea that a deterministic system can be driven to any final state in finite time given an external input^31^. This is commonly applied to linear, time invariant dynamic systems,

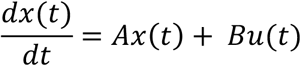

where *A* is an *NxN* matrix of state coefficients that describes how *N* molecule states, *x*(*t*), interact within the system and *B* is a matrix of input weights describing how external influences, *u*(*t*), impact the system. In general, a system is controllable if the controllability matrix,

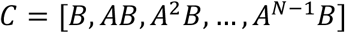

is full rank, *N*. This means that the system can be manipulated to reach any desired combination of states within all of state space following the defined input, *B*. In total, a controllability analysis identifies the key components of a system that must be manipulated to drive desired system outcomes^32^.

An example PPI network in Fig. 1a, is transformed into its state space matrix representation in Fig.1b. With the inclusion of two independent inputs (*u*_1_ and *u*_2_), the controllability matrix in Fig. 1c is full rank. Therefore, the system is fully controllable and it is possible to drive the protein concentrations to any desired state. Applying the idea of controllability to a cell at the onset of viral infection, a virus aims to control cellular functions (the system of proteins), promote virus replication tasks, and reach a final infected cell state. While it would be advantageous to interpret the infection from this control perspective, mathematical limits due to large system dimensions prevent the direct application of traditional controllability methods to PPI networks.

Advances in network theory have created alternative methods of network controllability evaluation which survey each node’s (protein’s) importance in the ability of an external set of inputs to fully control the network. Controllability classification is founded in “driver node” calculations: identifying the network components which must be manipulated for the system to be fully controlled (analogous to determining the non-zero elements of the *B* matrix in classic controllability). Without manipulation, driver nodes will remain unaffected by changes to the rest of the system, rendering the total system uncontrollable. A set of driver nodes (size *N_D_*) that is capable of controlling the total network is called a minimum input set (MIS). The MIS is not unique and the number of possible MISs scales exponentially with the size of the network^33^. After a primary MIS is calculated, two methods of controllability node classification can be used.

**Figure 1:**
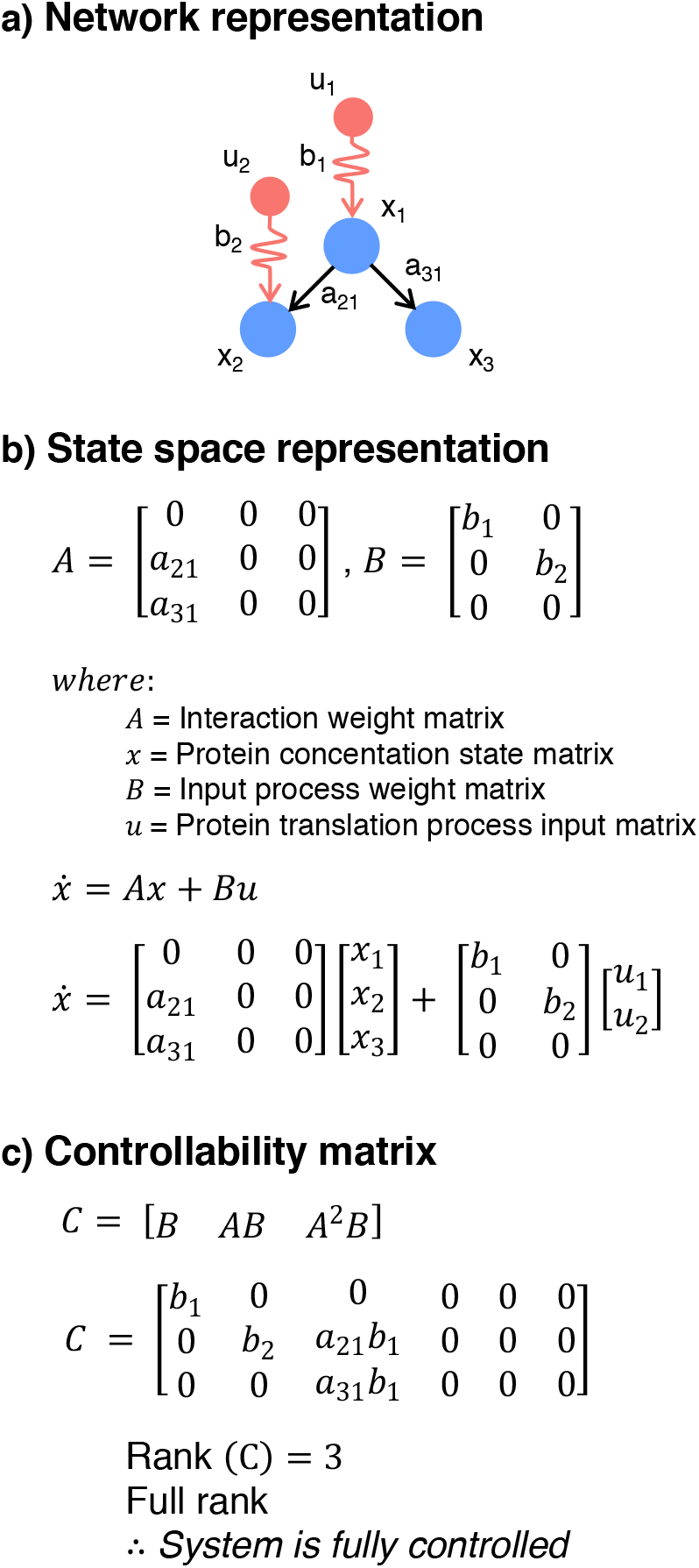
(a) An example protein-protein interaction network with three proteins and two protein translation process inputs. The state space representation (b) of the same network demonstrates that the change in state of a protein’s concentration is a function of its current state and an input process. A classic controllability analysis (c) demonstrates that this system is fully controllable and could, therefore, be driven to any possible state change in every protein.

In the first method by Liu et al.^34^, the mis is re-calculated (size *N_D_*′) after removing each node from the network. The node is then classified by its effect on the manipulation required to control the network, where an increase in the size of the MIS makes it more difficult to control the network and a decrease in the size of the MIS makes it easier to control the network. The absence of: an *indispensable node* increases the number of driver nodes (*N_D_*′ > *N_D_*), a *dispensable node* decreases the number of driver nodes (*N_D_*′ < *N_D_*), and a *neutral node* has no effect on the number of driver nodes (*N_D_*′ = *N_D_*). This method has previously been applied to many network types such as gene regulatory networks, food webs, citation networks, and PPI networks to better understand what drives the dynamics of each system^26,34^. While it is useful to observe the structural changes to the network in the absence of singular nodes, this method only considers one possible MIS. In a second classification method by Jia et al.^35^, a node is classified by its role across all possible MISs. A *critical node* is included in all possible MISs, an *intermittent node* is included in some possible MISs, and a *redundant node* is not included in any possible MISs. This method places each node in the broader context of all possible control configurations.

In total, this study aims to determine key host factors with regulatory roles specific to the influenza virus-infected cell state for the prediction of novel antiviral targets. We have completed a two-part controllability analysis of a host PPI network (HIN) and a hybrid network of human host PPI data combined with influenza A virus-host protein interaction data (VIN). The controllability characteristics of influenza virus interacting host proteins and driver proteins are compared to the characteristics of the total network. A set of 24 host factors that hold value topologically, in controllability, and functionally are identified as candidates for further study in drug development based on their specialized behavior during influenza infection.

## RESULTS

### Topology of the Host Interaction Network and Virus Integrated Network

The directed PPI network from Vinayagam et al^36^ was restricted to confident interactions (see Methods), creating a network containing 6,281 proteins and 31,079 interactions.

This network is referred to as the “Host Interaction Network” (HIN). Influenza A virus-host interactions from Watanabe et al^37^ were narrowed to 2,592 directed interactions between 11 influenza A virus (IAV) proteins (HA, M1, M2, NA, NP, NS1, NS2, PA, PB1, PB2, and PB1-F2 proteins) and 752 “IAV interacting proteins” preexisting in the HIN. After integration into the HIN, the network contains 6,292 proteins and 33,671 interactions. This network is referred to as the “Virus Integrated Network” (VIN).

Degree and betweenness calculations were completed for the HIN and VIN. As expected, the only proteins with altered degree after the addition of virus interactions to the network are the 752 IAV interacting proteins (Marked in blue in Fig. 2a). This shift is significant for the group of IAV interacting proteins as compared to all proteins in both the VIN (log scaled median of IAV interacting proteins: 1.04; log scaled median of all proteins: 0.70; student t-test of log scaled data p < 2.20×10^−16^) and the HIN (log scaled median of IAV interacting proteins: 0.85; log scaled median of all proteins: 0.70; Student t-test of log scaled data p: 5.97×10^−12^). The degree distributions of both networks are scale free (Fig. S1).

**Figure 2:**
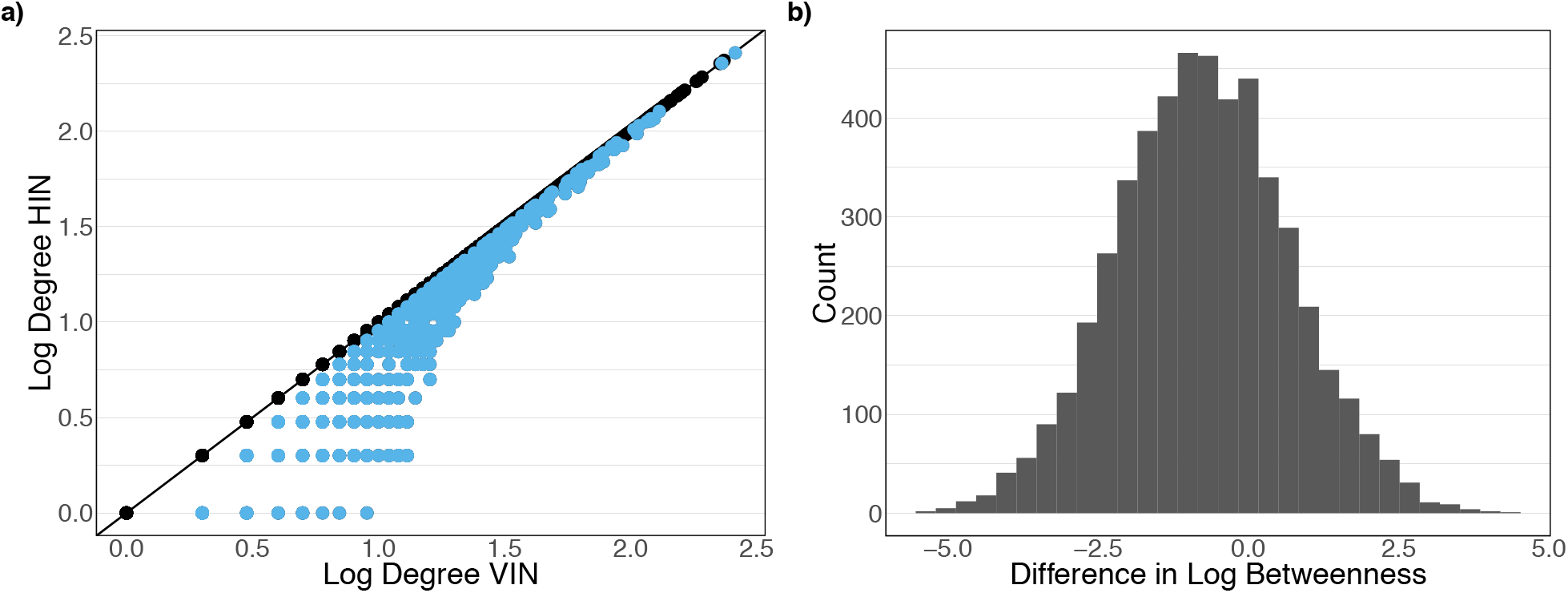
(a) Degree of the VIN vs degree of the HIN where the IAV interacting proteins are marked in blue. The degree distributions of the networks are scale free. (b) Difference in betweenness between the VIN and HINfor proteins which exhibit a difference greater than one.

Because betweenness is sensitive to the information flow through all proteins instead of only neighboring proteins, 2,735 proteins exhibit an increase in betweenness after the addition of IAV interactions. Of these proteins, 207 proteins’ log betweenness exhibits an increase of 2 or more in the VIN compared to the HIN (Fig. 2b). This suggests that the addition of IAV interactions has an effect on network topology that reaches over 3.5 times the number of host proteins that are directly interacting with IAV proteins. The betweenness shift in the group of IAV interacting host proteins is significant as compared to all proteins in both the VIN (Log scaled median of IAV interacting proteins: 3.23; Log scaled median of all proteins: 2.82; Student t-test of log scaled data p < 2.20×10^−16^) and the HIN (Log scaled median of IAV interacting proteins 3.22; Log scaled median of all proteins: 2.82; Student t-test of log scaled data p: 2.13×10^−15^). This is a result of being the limited protein set responsible for information flow from the viral proteins to the rest of the network.

### Driver proteins

Driver proteins (nodes) are the foundation of both types of controllability calculations,representing the protein set which must be manipulated for the system to be fully controlled. The proteins are identified through maximum matching algorithms^38^. The HIN and VIN both require *N_D_* = 2,463 driver proteins to achieve controllability, suggesting that the magnitude of network control is unchanged by the influence of the IAV interactions. However, the identity of driver proteins shifts slightly as the 11 viral proteins replace 11 host proteins within the primary MIS as drivers in the VIN. Table 1 lists their identities along with the shortest distance to an IAV protein in the network, degree, and betweenness. Of these 11 proteins, only five are directly interacting with IAV proteins. One of the remaining proteins is two steps (two interactions and one connecting protein) from any IAV protein, and the remaining five proteins are three steps from any IAV protein. The number of paths between viral proteins and these proteins are reflective of the number of paths between viral proteins and all host proteins (Fisher test p: 0.99). This supports the idea that viral interactions have lasting effects on the system’s control structure, affecting proteins that are multiple paths away.

**Table 1:** Identities of the proteins that are drivers in the HIN but not the VIN with the shortest number ofpaths to an Influenza A viral protein. Degree and betweenness of the proteins of the VIN is provided (with the values from the HIN in parenthesis). Only 45%o of these proteins are directly interacting with the viral proteins, demonstrating the cascade effect caused by the inclusion of viral interactions.

| Entrez ID | Gene Name | Shortest Distance to IAV Protein | Degree | Betweenness |
| --- | --- | --- | --- | --- |
| **10658** | CUGBP, Elav-Like Family Member 1 (CELF1) | 1 | 4 (4) | 81 (81) |
| **1969** | EPH Receptor A2 (EPHA2) | 1 | 14 (13) | 93 (0) |
| **6733** | SRSF Protein Kinase 2 (SRPK2) | 1 | 6 (2) | 6023 (6023) |
| **10318** | TNFAIP3 Interacting Protein 1 (TNIP1) | 1 | 7 (7) | 115 (115) |
| **2997** | Glycogen Synthase 1 (GYS1) | 3 | 4 (4) | 384 (384) |
| **10949** | Heterogeneous Nuclear Ribonucleoprotein A0 (HNRNPA0) | 2 | 9 (2) | 5 (0) |
| **64112** | Modulator of Apoptosis 1 (MOAP1) | 1 | 8 (8) | 6942 (6931) |
| **10419** | Protein Arginine Methyltransferase 5 (PRMT5) | 3 | 26 (17) | 6996 (4743) |
| **10262** | Splicing Factor 3b Subunit 4 (SF3B4) | 3 | 13 (7) | 82 (44) |
| **23321** | Tripartite Motif Containing 2 (TRIM2) | 3 | 2 (2) | 15 (15) |
| **81603** | Tripartite Motif Containing 8 (TRIM8) | 3 | 3 (3) | 0 (0) |

Lastly, analysis finds that 8.9% of all driver proteins are also IAV interacting proteins, meaning the intersection of the two protein groups of interest comprise only 3.5% of the total network. There is a significant increase in the betweenness of driver proteins depending on their status as IAV interacting or IAV non-interacting proteins (Fisher test p < 2.2×10^−16^) where there is no significant difference in degree of the same groups (Fisher test p: 0.7161). This is further evidence that the addition of virus interactions to the network magnifies information flow through the proteins most involved in controlling network behavior.

### Liu Controllability

Liu controllability was calculated (see Methods) for all proteins of the HIN and VIN (as shown in Table 2 with and without parentheses, respectively). The addition of IAV interactions to the network has no effect on the distribution of Liu classifications of host proteins, and consequently, the IAV Interacting proteins. Upon entry to the VIN, the 11 IAV proteins are classified as neutral, meaning their removal does not alter the number of driver proteins required to control the VIN (N_D_ = N_D_’). This reveals that the absence of singular proteins from the system is not enough to disturb the existing control structure under Liu controllability.

**Table 2:**
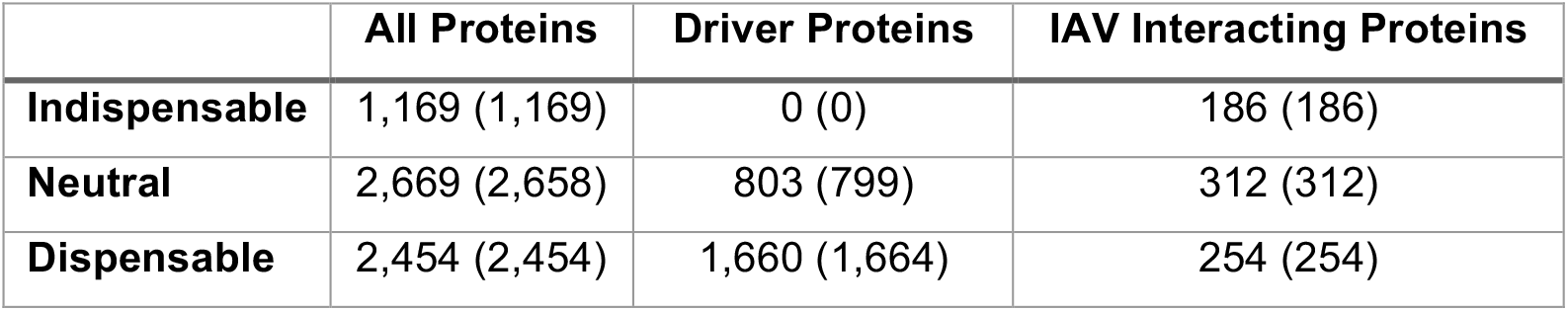
Liu types of all proteins, driver proteins, and virus interacting proteins in the VIN (HIN in parenthesis).

|  | **All Proteins** | **Driver Proteins** | **IAV Interacting Proteins** |
| --- | --- | --- | --- |
| **Indispensable** | 1,169 (1,169) | 0 (0) | 186 (186) |
| **Neutral** | 2,669 (2,658) | 803 (799) | 312 (312) |
| **Dispensable** | 2,454 (2,454) | 1,660 (1,664) | 254 (254) |

While none of the proteins change Liu classification between networks, the aforementioned replacement of 11 host driver proteins with viral proteins after the addition of virus interactions creates a small change in Liu type distribution for driver proteins. Of the displaced host proteins (deemed “Liu proteins”, found in Table 1), seven are neutral and four are redundant in the HIN, meaning that their removal from the network does not change the number of driver proteins and their removal reduces the number of driver proteins needed, respectively. Of the five Liu proteins that are both driver and IAV interacting proteins, four are neutral and one is redundant. The most notable change in degree and betweenness between the HIN and VIN is *PRMT5*, with an increase of 9 and 2,250, respectively. Overall, Liu controllability results suggest that the HIN is stable against potential changes in the control structure that could be caused by the addition of IAV interactions.

We developed an analysis to test if IAV is selectively targeting host proteins based on controllability characteristics. 10,000 random sets of 752 proteins (the number of IAV interacting proteins) were pulled from the host proteins of the VIN. Their Liu type distributions were plotted against the classification results of IAV interacting proteins, driver proteins, and all proteins in the VIN (Fig. 3a-c). The randomly sampled sets closely resemble all proteins of the network, not the true interacting protein set, suggesting that Liu controllability behavior of interacting proteins is not a coincidence of network construction (one-sided p = 0.51, 0.49, and 0.50 for indispensable, neutral, and dispensable, respectively). IAV interacting proteins tend to be indispensable compared to the percentage of all proteins that are indispensable (Fig. 3a). This suggests that viruses prefer to interact with proteins that are vital to cellular control. Driver proteins are very likely to be dispensable proteins compared to the percent of all proteins that are dispensable (Fig. 3c). Further, the mean and median log degree and betweenness of the randomly sampled protein sets is significantly lower than the same measurements of the true IAV interacting set (p < 2.2×10^−16^, 2.2×10^−16^, Fig. 4), signifying that virus interacting proteins are in positions of network significance. Overall, the Liu controllability results of IAV interacting proteins suggest that the virus may be selectively targeting host proteins based on controllability characteristics.

**Figure 3:**
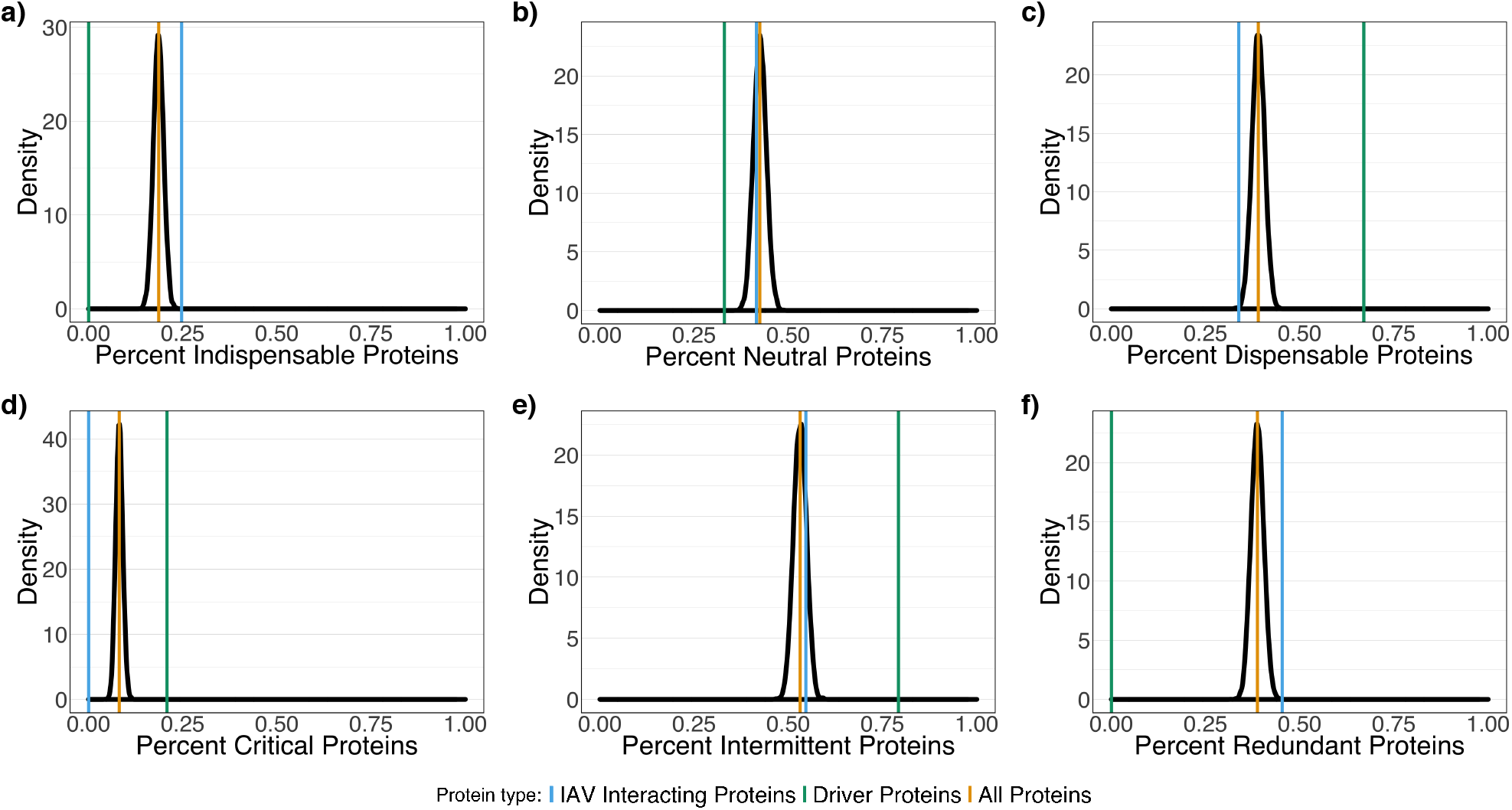
a-c) Density plots of distribution of Liu type for 10,000 random pulls of 752 proteins (number of virus interacting proteins in network). d-f Density plots of distribution of Jia type for 10,000 random pulls of 752 proteins (number of virus interacting proteins in network). Values for IAV interacting proteins (blue), driver proteins (green), and all proteins (gold) are pictured for all figures.

**Figure 4:**
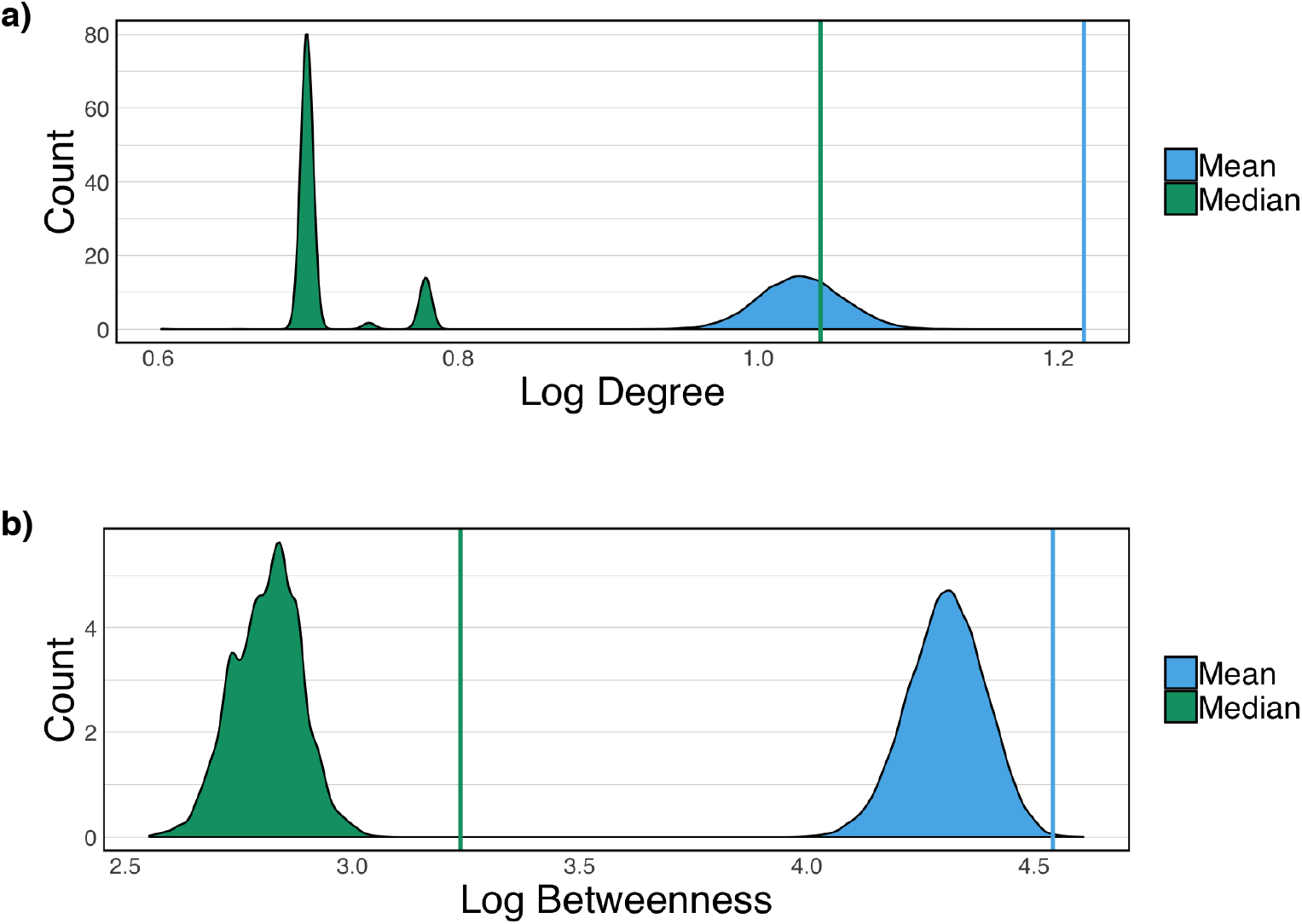
Density plots of a) mean (blue) and median (green) log degree of random IAV interacting protein sets and b) mean (blue) and median (green) log betweenness of random IAV interacting protein. Values for the true IAV interaction set shown as vertical lines, evidence that host proteins that directly interact with viral proteins are in positions of network significance.

### Jia Controllability

Jia controllability was calculated (see Methods) for all proteins of the HIN and VIN (as shown in Table 3 with and without parentheses, respectively). Unlike in Liu controllability, there is a small disturbance to Jia type distributions of host proteins after the addition of virus interactions. 24 host proteins shift from being classified as critical (a member of all MISs) to intermittent (a member of some MISs) proteins. Identities of these proteins (deemed “Jia proteins”) can be found in Table 4 along with the shortest distance to an IAV protein in the network and protein degree and betweenness. The two most notable changes in degree and betweenness between the HIN and VIN are EPH receptor A2 (*EPHA2*) with an increase of 1 and 93, respectively, and transferrin receptor (*TFRC*), with an increase of 3 and 164, respectively. All 24 Jia proteins are driver and IAV interacting proteins which, as mentioned, only comprises 3.5% of the total network. There are only two proteins (*EPHA2* and *HNRNPA0*) that are also members of the Liu protein set. 45% of IAV interacting proteins are never drivers, suggesting that they are always manipulated by neighboring host proteins within any possible control configuration. IAV interacting proteins are not enriched for driver proteins (Fisher test p: 0.14).

**Table 3:** Jia types of all proteins, driver proteins, and virus interacting proteins in the VIN (HIN in parenthesis)

|  | All Proteins | Driver Proteins | IAV Interacting Proteins |
| --- | --- | --- | --- |
| **Critical** | 512 (525) | 512 (525) | 0 (24) |
| **Intermittent** | 3,342 (3,318) | 1,951 (1,938) | 411 (387) |
| **Redundant** | 2,438 (2,438) | 0 (0) | 341 (341) |

Again, a randomized protein set was created to test if IAV may be selectively interacting with host proteins based on their controllability characteristics. 10,000 random sets of 752 proteins (the number of IAV interacting proteins) were sampled from the host proteins of the VIN. Their Jia type distributions were plotted against the classification results of IAV interacting proteins, driver proteins, and all proteins in the VIN (Fig. 3d-f). As with the Liu classification, the random sets closely resemble the total network (onesided p = 0.50, 0.51, and 0.50 for critical, intermittent, and redundant, respectively). While there are no redundant driver proteins by definition, driver proteins are more likely to be intermittent proteins than critical proteins (Fig. 3d-e), where more than 75% of all driver proteins are missing from at least one possible MIS. This means the majority of possible driver proteins are able to be controlled by a neighboring protein in at least one MIS. IAV interacting proteins tend to be redundant compared to the total number of proteins that are redundant (Fig. 3f). This suggests that viruses prefer to interact with proteins that are part of existing control structures to receive input from neighboring proteins.

**Figure 5:**
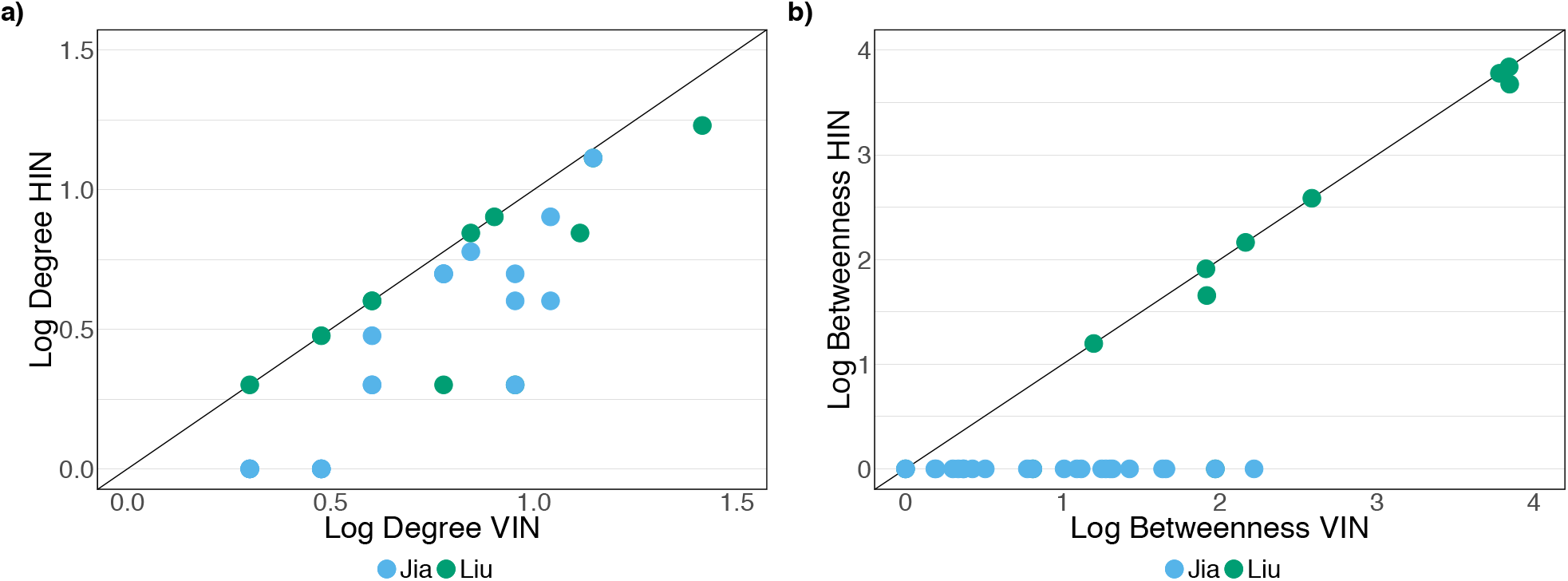
a) Degree and b) betweenness of Liu (blue) and Jia (green) protein sets between the HIN and VIN. While proteins identified in the Liu controllability analysis do not show significant deviation in degree or betweenness, proteins identified in the Jia controllability analysis show a shift in both measures after the addition of viral interactions.

Overall, Jia calculations identify a set of proteins for consideration that are more important within the VIN than the HIN. This is demonstrated through a comparison of degree and betweenness for the identified Liu and Jia driver sets in Fig. 5. Proteins identified in the Liu analysis show little deviation in both degree (Fig. 5a) and betweenness (Fig. 5b) measures after the addition of virus-host interactions to the network. In contrast, proteins identified in the Jia analysis show much larger deviations in degree (Fig. 5a) and betweenness (Fig. 5b) with all proteins having a betweenness of 0 in the HIN with an up to two log unit increase in the VIN (Table 4). Because the identified proteins were not responsible for information flow until the addition of virus-host interactions to the network, this suggests that the Jia protein set may identify key regulators of host immune response to infection.

**Table 4:** Identities of Jia Proteins (proteins that shift Jia classification between the HIN and VIN). All identified proteins are directly interacting with viral proteins. Degree and betweenness of the proteins of the VIN is provided (with the values from the HIN in parenthesis).

| **Entrez ID** | **Gene Name** | **Shortest Distance to IAV Protein** | **Degree** | **Betweenness** |
| --- | --- | --- | --- | --- |
| **56655** | DNA Polymerase Epsilon 4, Accessory Subunit (POLE4) | 1 | 2 (1) | 1 (0) |
| **30846** | EH Domain Containing 2 (EHD2) | 1 | 3 (1) | 1 (0) |
| **1969** | EPH Receptor A2 (EPHA2) | 1 | 14 (13) | 93 (0) |
| **2665** | GDP Dissociation Inhibitor 2 (GDI2) | 1 | 3 (1) | 2 (0) |
| **51552** | RAB14, Member RAS Oncogene Family (RAB14) | 1 | 2 (1) | 1 (0) |
| **2091** | Fibrillarin (FBL) | 1 | 9 (4) | 19 (0) |
| **10949** | Heterogeneous Nuclear Ribonucleoprotein A0 (HNRNPA0) | 1 | 9 (2) | 5 (0) |
| **3032** | Hydroxyacyl-CoA Dehydrogenase/3-Ketoacyl-CoA Thiolase/Enoyl-CoA Hydratase (Trifunctional Protein), Beta Subunit (HADHB) | 1 | 9 (5) | 26 (0) |
| **3419** | Isocitrate Dehydrogenase 3 (NAD(+)) Alpha (IDH3A) | 1 | 3 (1) | 2 (0) |
| **4191** | Malate Dehydrogenase 2 (MDH2) | 1 | 3(1) | 1 (0) |
| **64949** | Mitochondrial Ribosomal Protein S26 (MRPS26) | 1 | 2 (1) | 0 (0) |
| **9180** | Oncostatin M Receptor (OSMR) | 1 | 6 (5) | 18 (0) |
| **5052** | Peroxiredoxin 1 (PRDX1) | 1 | 11 (4) | 44 (0) |
| **5213** | Phosphofructokinase, Muscle (PFKM) | 1 | 6 (5) | 17 (0) |
| **26227** | Phosphoglycerate Dehydrogenase (PHGDH) | 1 | 4 (2) | 9 (0) |
| **5817** | Poliovirus Receptor (PVR) | 1 | 7 (6) | 42 (0) |
| **5686** | Proteasome Subunit Alpha 5 (PSMA5) | 1 | 6 (5) | 11 (0) |
| **5464** | Pyrophosphatase (Inorganic) 1 (PPA1) | 1 | 6 (5) | 5 (0) |
| **113174** | Serum Amyloid A Like 1 (SAAL1) | 1 | 2 (1) | 1 (0) |
| **6745** | Signal Sequence Receptor Subunit 1 (SSR1) | 1 | 4 (2) | 12 (0) |
| **7037** | Transferrin Receptor (TFRC) | 1 | 11 (8) | 164 (0) |
| **8834** | Transmembrane Protein 11 (TMEM11) | 1 | 4 (3) | 20 (0) |
| **30000** | Transportin 2 (TNPO2) | 1 | 2 (1) | 1 (0) |
| 7407 | Valyl-Trna Synthetase (VARS) | 1 | 3 (1) | 0 (0) |

### Validation of controllability significant host factors

All proteins were checked against 6 siRNA screens for host factors involved in influenza replication (Brass et al^39^, Hao et al^40^, Karlas et al^41^, König et al^42^, Shapira et al^43^, and Watanabe et al^37^), grouped by both Liu and Jia controllability classifications. Less than 5% of all classifications of both types are validated by any of the 6 screens (Fig. 6), suggesting that no controllability classification is more enriched for host factors than another. This is likely due to the low agreement observed across siRNA studies^44^. However, the driver proteins that change Liu and Jia classification have higher hit rates in siRNA screens, with 2 of 11 changing MIS proteins (*SF3B4, SRPK2*, 18% validation) and 5 of 24 Jia-identified proteins (*OSMR, PPA1, PSMA5, POLE4, GDI2*, 21% validation), though neither are statistically significant results (Fisher p-values of 0.685 and 0.252, respectively).

**Figure 6:**
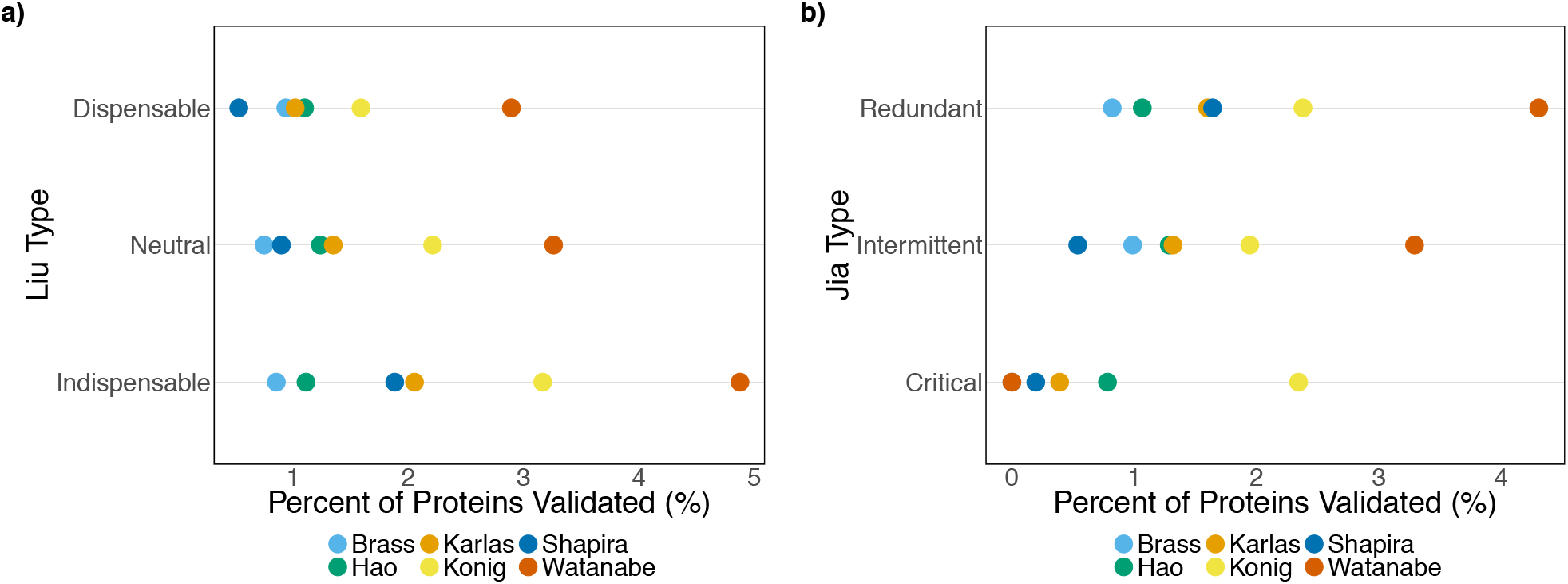
Percent of each a) Liu classification type and b) Jia classification type confirmed in 6 siRNA screens (Brass, Karlas, Shapira, Hao, Konig, Watanabe). None of the 6 possible classifications are more than 5% validated in the screenings, suggesting that experimental findings do not favor certain protein controllability types.

An analysis of both protein sets of interest was performed using Ingenuity Pathway Analysis (IPA)^45^. The network created for the Liu protein set identified cellular compromise, cell death, and cell cycle functions. The network created for the Jia protein set identified protein synthesis functions, all centered around NF-kB. The Jia network notably recognizes six proteins (*EPHA2, FBL, PFKM, PSMA5, SSR1*, and *TFRC*) for their involvement in the infection of cells (p:9.58×10^−4^). Four proteins in the Liu network (*CELF1, SF384, SRPK2*, and *HNRNPA0*, the last of which appears in both protein sets) were identified for their involvement in mRNA processing (p-value: 3.33×10^−6^).

Lastly, Interferome v2.01^46^ was used to determine if the 11 Liu proteins and 24 Jia proteins are interferon regulated genes (IRGs). All 11 Liu proteins are identified as IRGs and exhibit a 2-fold change in expression when treated with interferon in at least one experimental dataset. 20 of 24 Jia proteins are identified as IRGs and exhibit a 2-fold change in expression in at least one experimental dataset. 6 Jia proteins are identified in more than 10 studies. In particular, *HNRNPA0* and *PPA1* are significantly down regulated in 20 and 63 datasets, respectively. These results point toward the involvement of the predicted protein subsets in immune response events.

## DISCUSSION

In total, this two-part network controllability analysis for a host protein-protein interaction network (HIN) and an integrated influenza virus-host protein-protein interaction network (VIN) aims to enhance the prediction of antiviral drug targets for influenza A virus. While Liu controllability methods have previously been applied to study PPI networks^26^, past analysis focuses only on the classification of virus interacting proteins and does not evaluate before and after the addition of virus-host interactions to the network. A Jia controllability analysis has never been applied to PPI networks. The unique construction of the VIN includes experimentally-derived virus-host interaction data^37^ which represents opportunities for the virus to manipulate host intracellular machinery using protein-protein interactions. Here, analysis of the transition between the healthy and infected network states and further investigation of virus interacting and driver proteins has identified 24 proteins as regulatory markers of the infected state. This protein set is noted for its characteristics in topology, controllability, and functional roles within the infected cell: results that are summarized in Table 5. Our workflow observes both the effect of structural changes to the network in the case of potential protein knock outs, as well as each protein’s role in all MISs, representing all possible ways of controlling the system. In combination, network approach and results provide deeper understanding of how changes to cell behavior at the onset of infection are able to occur through the work of a small set of viral proteins. Through understanding the system in this way, we present the possibility to “outsmart” viral attack by dismantling the control structure which allows the viral infection to take hold.

**Table 5:** Summary of results for proteins identified in the Jia controllability analysis.

| **Quality** | **Frequency in Jia Protein Set** |
| --- | --- |
| Driver protein | 100% |
| IAV interacting protein | 100% |
| Identified in Liu protein set | 8% |
| Validated in at least one siRNA screen | 21% |
| Cell infection – functional enrichment | 25% |
| mRNA processing – functional enrichment | 17% |
| Interferon regulating gene | 83% |

A network representation of the cellular environment demonstrates that the effects of infection (represented by the addition of virus-host interactions) cascade through the system, demonstrated by the alteration of basic topology measures. The betweenness shift between the two networks, particularly in IAV interacting proteins, supplies evidence that the topological effect of viral infection is wide reaching (Tables 1 and 4). Further, a comparison of driver protein betweenness for those that are also IAV interacting proteins in comparison to those that are not shows a significant difference. By definition, driver proteins that are IAV interacting are not receiving control influence from viral proteins and require additional external influence to achieve network control. However, the increased betweenness of proteins that are both driver and IAV interacting proteins suggests that this group is still of great importance to information flow through the network. This is one example where differences in network topology measures can emphasize the importance of select proteins that are overlooked by controllability principles.

Controllability analyses confirm that IAV interacting proteins are in positions of significance for both types of classification. The increased population of indispensable IAV interacting proteins (Liu controllability: *N_D_*′ > *N6*, Fig. 3a) compared to what would be expected by random chance suggests that it would be more difficult for an outside influence (such as viral infection) to control the network in the absence of IAV interacting proteins opposed to a randomly selected protein. This is logical as IAV interacting proteins act as the connection between viral proteins and the host network where control is initiated. The increased population of redundant IAV interacting proteins (Jia controllability: never a driver protein, Fig. 3f) when compared to the random expectation indicates that more IAV interacting proteins are always being manipulated internally than would be expected by chance. This means that they are fully incorporated into the control structure of the VIN. From these two results, one can conclude that IAV interacting proteins contribute to both the “gate” (the ease of entering the system) and the “heart” (the proteins responsible for propagating control through the system) of the network control structure during infection. These findings support the idea that viruses are likely to interact with proteins which offer an advantage to total network control.

Similarly, both sets of controllability results demonstrate that driver proteins play interesting roles in the network control structure. The large population of dispensable driver proteins (Liu controllability: *N_D_*′ < *N_D_*, Table 2) signifies that the majority of driver proteins are making it more difficult to control the network by requiring more external inputs to control system behavior. In their absence, the number of driver proteins would decrease and it would theoretically be easier for a viral attack to compromise the network control structure. As such, a possible strategy for drug development could be to protect these proteins from drastic changes to abundance. Over 75% of driver proteins are classified as intermittent (Jia controllability: sometimes a driver protein, Table 3), meaning there is at least one MIS where these driver proteins are not drivers, and receive control influence through internal propagation. This lends itself to the idea of viral escape routes: under pressure, virus proteins could utilize alternative pathways to maintain system control and reach the goal of hijacking cellular function.

The method of controllability implementation used identifies protein sets of interest through changes to classification between the HIN and VIN. Unfortunately, Liu classification methods do not detect a change between the two networks in this study. As it is a measure of the robustness of the network to structural changes in the absence of each protein, this suggests that the HIN upholds its typical control structure during IAV infection. This could be a consequence of the interaction data used or it may be that the strategy applied here cannot distinguish between the behavior of healthy and diseased states. Knowing the extent of changes to cell behavior within immune response pathways^47–49^, apoptosis signaling^50,51^, and transcriptional processes^52–54^ during infection, the IAV infected cell can be interpreted as a different system. The failure to see this distinction may be a shortcoming of the Liu controllability calculation, especially knowing that the 11 Liu proteins are not unique due to the method’s use of a single MIS. Overall, the Liu analysis should be applied to additional virus-host networks in the fashion described within this study to further evaluate the method.

The 24 proteins identified by the Jia controllability analysis show promise as indicators of regulatory roles specific to the infected state. All Jia proteins are IAV interacting and driver proteins, a high distinction which demonstrates a significant importance to network information flow marked by significantly higher betweenness in the VIN than even driver proteins that are not IAV interacting. Additionally, all Jia proteins have no importance to network flow in the HIN (betweenness = 0) (Table 4), suggesting their role in network structure “turns on” after the onset of infection. It is noteworthy that *PRDX1* has been implicated in respiratory syncytial virus (RSV)^55^, a lower respiratory tract infection that is often associated with influenza virus^56^. Though the number of Jia proteins identified in existing siRNA screening data is not statistically significant, it should be noted that siRNA screens cover only the partial genome. As such, this type of analysis could be used to direct future experimental studies to save time, money, and effort. IPA analysis reveals that some of the identified proteins hold roles in mRNA processing, an integral part of the influenza virus’ ability to spread through processing its own RNA using host machinery^57^. The Jia protein network is centered around NF-kB, which is implicated in host immunity with evidence that the virus directly inhibits NF-kB activity^58,59^. The interferon regulating roles of proteins in a high number of both identified sets (all 11 changing MIS proteins and 20 of 24 Jia-identified proteins) speak to their responsibility in controlling infection. *PPA1* appears as downregulated in 63 studies and *HNRNPA0* appears as downregulated in 20 studies when treated with interferon compared to a control, solidifying their involvement in the host immune response. In total, this evidence suggests that controllability analyses hold power as predictors for important regulators of the host response to influenza infection and, therefore, hold power for drug target prediction.

Existing influenza virus studies using PPI networks require additional data such as differentially expressed gene information^60^ or protein context^27^ to construct host response networks. Alternative methods such as DeltaNet^61,62^ and ProTINA^63^ utilize gene transcription profiles to infer protein drug targets, but rely on the accurate deduction of gene regulatory networks. More recent PPI studies have used network growing functions such as GeneMANIA, STRING, and IPA^64^ to predict IAV host factors and studied infected cell systems through the integration of screening data with network methods^30,65^. Approaches incorporating time course data into network analysis have also been explored^66^. While these methods (which include basic network metrics such as degree and betweenness of PPI networks) have been successful at identifying disease host factors and in drug target development in the existing body of work, this dual controllability study offers a novel, in-depth analysis of the role of individual proteins in the context of total system function and how possible changes to the system can be interpreted.

Lastly, though this study has used experimental data from Influenza A studies, this analysis can be used to improve the prediction of drug targets for any pathogen-host interaction given available protein interaction data because of the generality of the method. The limits of these methods lie in limited availability of large-scale, dependable databases of protein-protein interactions. Foundational maximum matching algorithms for the calculation of driver proteins must be performed with directed networks. While larger directed networks than the network from Vinayagam et al.^36^ are available^67^, the network used here contains only experimentally derived data opposed to computationally predicted interactions, assuring biological confidence in the results within this study. A Liu controllability analysis of the computationally predicted network presented in Uhart et al.^67^ finds that 29% of proteins are categorized as indispensable where approximately 20% of proteins in the Vinayagam network are classified as the same, though there is 89% overlap in directed edges between the two networks. This suggests that methods for predicting protein interactions may over represent these key proteins within the analysis, even in combination with experimental results. However, larger networks will move towards a more complete analysis of infected cell behavior and possibly reveal further proteins of interest. Therefore, the future of this field depends on continued establishment of large, confident, directed PPI networks.

## METHODS

### Protein-protein interaction network

The host protein-protein interaction network used was downloaded from Vinayagam et al^36^. A confidence level cutoff of 0.7 was used, creating the HIN. Influenza A virus-host interactions from Watanabe et al^37^ were narrowed to interactions which contained host proteins already found in host network interactions to avoid skewing degree and betweenness network metrics. These interactions were directly integrated into the host network, creating the VIN. All analysis was completed in R 3.4.3 using the igraph package.

### Liu classification

Calculations for Liu classification were adopted from Liu et al^34^. For a network of *n* nodes, a set of driver nodes for the bipartite representation of the network, *N_D_*, is found using a maximum matching algorithm such as Hopcroft-Karp^38^. Each node of the network is iteratively removed (*N’* = *N* – 1) and maximum matching, *N_D_*′, is reevaluated. Nodes are classified as indispensable (*N_D_*′ > *N_D_*), neutral (*N_D_*′ = *N_D_*), or dispensable *N_D_*′ < *N_D_*).

### Jia classification

Calculations for Jia classification were adopted from Jia et al^35^. For a network of *n* nodes, a set of driver nodes for the bipartite representation of the network, *N_D_*, is found using a maximum matching algorithm such as Hopcroft-Karp^38^. For all *N_D_*, control adjacent nodes were identified iteratively and an input graph was created as dictated in Zhang et al^68^. The input graph was used to classify nodes as critical (in all minimum input sets), neutral (in some minimum input sets), or redundant (in no minimum input sets).

## ACKNOWLEDGEMENTS

Thank you to the Department of Chemical and Petroleum Engineering at the University of Pittsburgh for funding this research. Thank you to the Systems Biology Institute, Tokyo, for expertise and computational training. Thank you to the Howard Hughes Medical Institute for supporting this project through the James H. Gilliam Fellowships for Advanced Study program.

## AUTHOR CONTRIBUTIONS

Emily E. Ackerman assisted in conceptualization of the study, designed the study, performed all computational experiments, and wrote the manuscript. Jason E. Shoemaker conceptualized and funded the study. John F. Alcorn assisted in conceptualization of the study and advised on relevant virology and immunology. Takeshi Hase provided computational training.

---

**Fig. S1:**
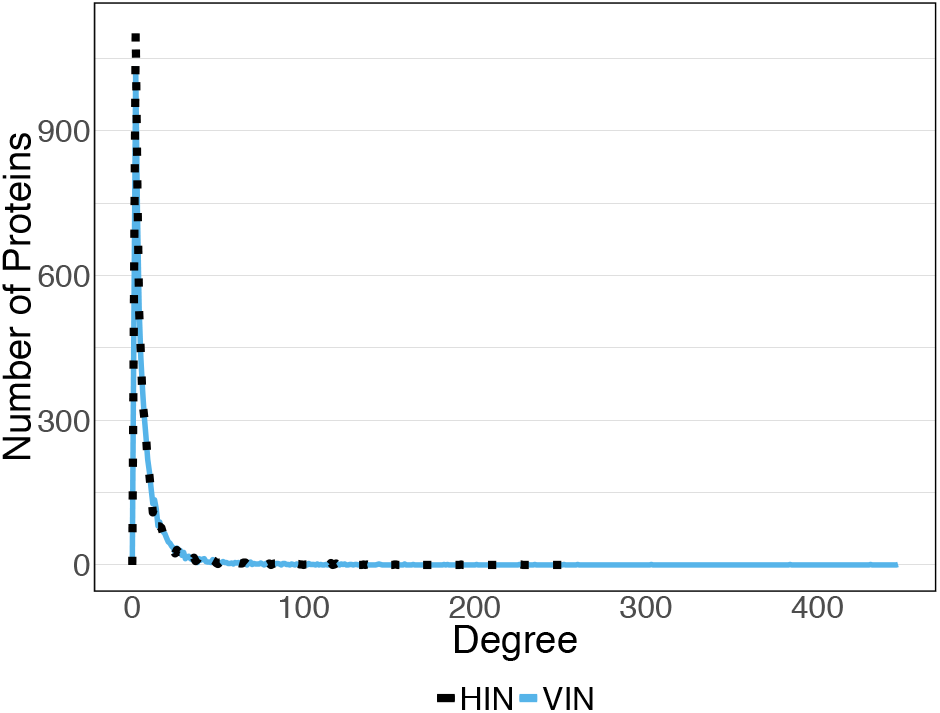
Degree distribution of network with IAV interactions (blue solid) and without IAV interactions (dotted black) show that both networks demonstrate scale free topology

## CITATIONS

1. Rask-Andersen, M., Almén, M. S. & Schiöth, H. B. Trends in the exploitation of novel drug targets. Nat. Rev. Drug Discov. (2011). doi:10.1038/nrd3478

2. Klipp, E. & Liebermeister, W. Mathematical modeling of intracellular signaling pathways. BMC Neuroscience 7, (2006).

3. Schoeberl, B., Eichler-Jonsson, C., Gilles, E. D. & Muüller, G. Computational modeling of the dynamics of the MAP kinase cascade activated by surface and internalized EGF receptors. Nat. Biotechnol. 20, 370–375 (2002).

4. Aldridge, B. B., Burke, J. M., Lauffenburger, D. A. & Sorger, P. K. Physicochemical modelling of cell signalling pathways. Nature Cell Biology 8, 1195–1203 (2006).

5. Cho, D.-Y., Kim, Y.-A. & Przytycka, T. M. Chapter 5: Network Biology Approach to Complex Diseases. PLoS Comput. Biol. 8, e1002820 (2012).

6. Freeman, L. C. A Set of Measures of Centrality Based on Betweenness. Sociometry 40, 35–41 (1977).

7. Zhu, M. et al. The analysis of the drug-targets based on the topological properties in the human protein-protein interaction network. J. Drug Target. 17, 524–532 (2009).

8. Vinayagam, A. et al. Integrating protein-protein interaction networks with phenotypes reveals signs of interactions. Nat. Methods 11, 94–99 (2014).

9. He, X. & Zhang, J. Why do hubs tend to be essential in protein networks? PLoS Genet. (2006). doi:10.1371/journal.pgen.0020088

10. Lopes, T. J. S., Shoemaker, J. E., Matsuoka, Y., Kawaoka, Y. & Kitano, H. Identifying problematic drugs based on the characteristics of their targets. Front. Pharmacol. 6, (2015).

11. Barabási, A.-L., Gulbahce, N. & Loscalzo, J. Network medicine: a network-based approach to human disease. Nat. Rev. Genet. 12, 56–68 (2011).

12. Jonsson, P. F. & Bates, P. A. Global topological features of cancer proteins in the human interactome. Bioinformatics 22, 2291–2297 (2006).

13. Hase, T., Tanaka, H., Suzuki, Y., Nakagawa, S. & Kitano, H. Structure of protein interaction networks and their implications on drug design. PLoS Comput. Biol. 5, (2009).

14. Mani, K. M. et al. A systems biology approach to prediction of oncogenes and molecular perturbation targets in B-cell lymphomas. Mol. Syst. Biol. (2008). doi:10.1038/msb.2008.2

15. Mine, K. L. et al. Gene network reconstruction reveals cell cycle and antiviral genes as major drivers of cervical cancer. Nat. Commun. (2013). doi:10.1038/ncomms2693

16. Mitchell, H. D. et al. A Network Integration Approach to Predict Conserved Regulators Related to Pathogenicity of Influenza and SARS-CoV Respiratory Viruses. PLoS One (2013). doi:10.1371/journal.pone.0069374

17. Gandhi, T. K. B. et al. Analysis of the human protein interactome and comparison with yeast, worm and fly interaction datasets. Nat. Genet. (2006). doi:10.1038/ng1747

18. Arrell, D. K. & Terzic, A. Network systems biology for drug discovery. Clinical Pharmacology and Therapeutics 88, 120–125 (2010).

19. Pujol, A., Mosca, R., Farrés, J. & Aloy, P. Unveiling the role of network and systems biology in drug discovery. Trends in Pharmacological Sciences 31, 115–123 (2010).

20. Germain, M.-A. et al. Elucidating novel hepatitis C virus-host interactions using combined mass spectrometry and functional genomics approaches. Mol. Cell. Proteomics 13, 184–203 (2014).

21. de Chassey, B. et al. Hepatitis C virus infection protein network. Mol. Syst. Biol. 4, 1–12 (2008).

22. Moni, M. A. & Liò, P. Network-based analysis of comorbidities risk during an infection: SARS and HIV case studies. BMC Bioinformatics (2014). doi:10.1186/1471-2105-15-333

23. Murali, T. M., Dyer, M. D., Badger, D., Tyler, B. M. & Katze, M. G. Network-based prediction and analysis of HIV dependency factors. PLoS Comput. Biol. (2011). doi:10.1371/journal.pcbi.1002164

24. Ptak, R. G. et al. Short Communication: Cataloguing the HIV Type 1 Human Protein Interaction Network. AIDS Res. Hum. Retroviruses (2008). doi:10.1089/aid.2008.0113

25. Shityakov, S., Dandekar, T. & Förster, C. Gene expression profiles and protein-protein interaction network analysis in AIDS patients with HIV-associated encephalitis and dementia. HIV/AIDS-Res. Palliat. Care (2015). doi:10.2147/HIV.S88438

26. Vinayagam, A. et al. Controllability analysis of the directed human protein interaction network identifies disease genes and drug targets. Proc. Natl. Acad. Sci. 113, 4976–4981 (2016).

27. Schaefer, M. H. et al. Adding protein context to the human protein-protein interaction network to reveal meaningful interactions. PLoS Comput. Biol. 9, e1002860 (2013).

28. Shoemaker, J. E. et al. Integrated network analysis reveals a novel role for the cell cycle in 2009 pandemic influenza virus-induced inflammation in macaque lungs. BMC Syst. Biol. 6, 117 (2012).

29. Korth, M. J., Tchitchek, N., Benecke, A. G. & Katze, M. G. Systems approaches to influenza-virus host interactions and the pathogenesis of highly virulent and pandemic viruses. Semin. Immunol. 1–12 (2012). doi:10.1016/j.smim.2012.11.001

30. Tripathi, S. et al. Meta-and Orthogonal Integration of Influenza ‘oMICs’ Data Defines a Role for UBR4 in Virus Budding. Cell Host Microbe 18, 723–735 (2015).

31. Lin, C. T. Structural Controllability. IEEE Trans. Automat. Contr. 19, 201–208 (1974).

32. Wuchty, S. Controllability in protein interaction networks. Proc. Natl. Acad. Sci. (2014). doi:10.1073/pnas.1311231111

33. Jia, T. & Barabási, A. L. Control capacity and a random sampling method in exploring controllability of complex networks. Sci. Rep. 3, (2013).

34. Liu, Y. Y., Slotine, J. J. & Barabási, A. L. Controllability of complex networks. Nature 473, 167–173 (2011).

35. Jia, Tao; Liu, Yang-Yu; Csòka, Endre; Pòsfai, Márton; Slotine, Jean-Jacques; Barabáasi, A.-L. Emergence of bimodality in controlling complex networks. Nat. Commun. (2013). doi:10.1038/ncomms3002

36. Vinayagam, A. et al. A directed protein interaction network for investigating intracellular signal transduction. Sci. Signal. (2011). doi:10.1126/scisignal.2001699

37. Watanabe, T. et al. Influenza virus-host interactome screen as a platform for antiviral drug development. Cell Host Microbe 16, 795–805 (2014).

38. Hopcroft, J. E. & Karp, R. M. An $n^{5/2} $ Algorithm for Maximum Matchings in Bipartite Graphs. SIAM J. Comput. (1973). doi:10.1137/0202019

39. Brass, A. L. et al. The IFITM proteins mediate cellular resistance to influenza A H1N1 virus, West Nile virus, and dengue virus. Cell 139, 1243–1254 (2009).

40. Hao, L. et al. Drosophila RNAi screen identifies host genes important for influenza virus replication. Nature 454, 890–893 (2008).

41. Karlas, A. et al. Genome-wide RNAi screen identifies human host factors crucial for influenza virus replication. Nature 463, 818–22 (2010).

42. König, R. et al. Human host factors required for influenza virus replication. Nature 463, 813–7 (2010).

43. Shapira, S. D. et al. A physical and regulatory map of host-influenza interactions reveals pathways in H1N1 infection. Cell 139, 1255–1267 (2009).

44. Hao, L. et al. Limited agreement of independent RNAi screens for virus-required host genes owes more to false-negative than false-positive factors. PLoS Comput. Biol. 9, e1003235 (2013).

45. Krämer, A., Green, J., Pollard, J. & Tugendreich, S. Causal analysis approaches in ingenuity pathway analysis. Bioinformatics 30, 523–530 (2014).

46. Samarajiwa, S. A., Forster, S., Auchettl, K. & Hertzog, P. J. INTERFEROME: The database of interferon regulated genes. Nucleic Acids Res. (2009). doi:10.1093/nar/gkn732

47. Koyama, S., Ishii, K. J., Coban, C. & Akira, S. Innate immune response to viral infection. Cytokine 43, 336–341 (2008).

48. Thompson, M. R., Kaminski, J. J., Kurt-Jones, E. A. & Fitzgerald, K. A. Pattern recognition receptors and the innate immune response to viral infection. Viruses (2011). doi:10.3390/v3060920

49. Iwasaki, A. & Medzhitov, R. Toll-like receptor control of the adaptive immune responses. Nature Immunology (2004). doi:10.1038/ni1112

50. Barber, G. N. Host defense, viruses and apoptosis. Cell Death and Differentiation (2001). doi:10.1038/sj.cdd.4400823

51. Thomson, B. J. Viruses and apoptosis. International Journal of Experimental Pathology (2001). doi:10.1046/j.1365-2613.2001.00195.x

52. Gale Jr, M., Tan, S.-L. & Katze, M. G. Translational control of viral gene expression in eukaryotes. Microbiol. Mol. Biol. Rev. (2000). doi:10.1128/mmbr.64.2.239-280.2000

53. Sonenberg, N. & Hinnebusch, A. G. Regulation of Translation Initiation in Eukaryotes: Mechanisms and Biological Targets. Cell (2009). doi:10.1016/j.cell.2009.01.042

54. Walsh, D., Mathews, M. B. & Mohr, I. Tinkering with translation: Protein synthesis in virus-infected cells. Cold Spring Harb. Perspect. Biol. (2013). doi:10.1101/cshperspect.a012351

55. Pavia, A. T. Viral infections of the lower respiratory tract: Old viruses, new viruses, and the role of diagnosis. Clin. Infect. Dis. (2011). doi:10.1093/cid/cir043

56. Jamaluddin, M. et al. Role of Peroxiredoxin 1 and Peroxiredoxin 4 in Protection of Respiratory Syncytial Virus-Induced Cysteinyl Oxidation of Nuclear Cytoskeletal Proteins. J. Virol. (2010). doi:10.1128/JVI.01005-10

57. Dubois, J., Terrier, O. & Rosa-Calatrava, M. Influenza viruses and mRNA splicing: Doing more with less. mBio (2014). doi:10.1128/mBio.00070-14

58. Kumar, N., Xin, Z.-T., Liang, Y., Ly, H. & Liang, Y. NF-kappaB signaling differentially regulates influenza virus RNA synthesis. J. Virol. 82, 9880–9 (2008).

59. Ludwig, S. & Planz, O. Influenza viruses and the NF-κB signaling pathway-Towards a novel concept of antiviral therapy. Biological Chemistry 389, 1307–1312 (2008).

60. Shoemaker, J. E. et al. Integrated network analysis reveals a novel role for the cell cycle in 2009 pandemic influenza virus-induced inflammation in macaque lungs. BMC Syst. Biol. 6, (2012).

61. Noh, H. & Gunawan, R. Inferring gene targets of drugs and chemical compounds from gene expression profiles. Bioinformatics (2016). doi:10.1093/bioinformatics/btw148

62. Noh, H., Ziyi, H. & Gunawan, R. Inferring Causal Gene Targets from Time Course Expression Data. IFAC-PapersOnLine (2016). doi:10.1016/j.ifacol.2016.12.151

63. Noh, H., Shoemaker, J. E. & Gunawan, R. Network perturbation analysis of gene transcriptional profiles reveals protein targets and mechanism of action of drugs and influenza A viral infection. Nucleic Acids Res. (2018). doi:10.1093/nar/gkx1314

64. Taye, B. et al. Benchmarking selected computational gene network growing tools in context of virus-host interactions. Sci. Rep. 7, 5805 (2017).

65. Heaton, N. S. et al. Targeting Viral Proteostasis Limits Influenza Virus, HIV, and Dengue Virus Infection. Immunity 44, 46–58 (2016).

66. Jain, S., Arrais, J., Venkatachari, N. J., Ayyavoo, V. & Bar-Joseph, Z. Reconstructing the temporal progression of HIV-1 immune response pathways. Bioinformatics (2016). doi:10.1093/bioinformatics/btw254

67. Uhart, M., Flores, G. & Bustos, D. M. Controllability of protein-protein interaction phosphorylation-based networks: Participation of the hub 14-3-3 protein family. Sci. Rep. (2016). doi:10.1038/srep26234

68. Zhang, X., Lv, T. & Pu, Y. Input graph: The hidden geometry in controlling complex networks. Sci. Rep. (2016). doi:10.1038/srep38209

